# Male-dominated courtship in an unexpectedly late-breeding Andean Lapwing (*Vanellus resplendens*) population

**DOI:** 10.1101/2022.10.25.513697

**Authors:** Guillaume Dillenseger, Andreas Rimoldi, Santiago Barreto, L. Mauricio Ugarte-Lewis, Vojtěch Kubelka

## Abstract

Sexual behavior, namely courtship and mating strategies, remain unexplored for numerous tropical birds. Andean Lapwing (*Vanellus resplendens*) is a shorebird inhabiting High Andes, and its reproductive biology is poorly known. Here, we performed standardized 49 observations of 20 pairs of Andean Lapwings at Laguna de Salinas, Peru (4315 m a.s.l.) during 2021–2022 breeding season. Lapwings in this studied population show social monogamy, a defined breeding season apparently triggered by rainfall, starting later than what is known according to the literature, and male-biased investment in courtship and territorial defense. These observations increase our knowledge on this understudied Neotropical species breeding at high elevations.

**RESUMEN:** El comportamiento sexual, es decir, las estrategias de cortejo y apareamiento, permanecen sin explorarse para numerosas aves tropicales. El Avefría Andina (*Vanellus resplendens*) es un ave playera que habita los altos Andes, sin embargo, su biología reproductiva es poco conocida. Aquí, realizamos 49 observaciones estandarizadas de 20 parejas de avefrías andinas en Laguna de Salinas, Perú (4315 m.s.n.m.) durante la temporada 2021–2022. Las avefrías en esta población estudiada muestran monogamia social, una temporada reproductiva bien definida provocada por las precipitaciones, comenzando más tarde de lo que es descrito en la literatura, y una inversión predispuesta de machos en el cortejo y la defensa territorial. Estas observaciones aumentan nuestro conocimiento sobre esta especie Neotropical poco estudiada que se reproduce a gran altitud.

---

Sexual behaviors are among the most variable behaviors in animal kingdom, also predominantly determining reproductive success of involved individuals (e.g., Parish & Coulson, 1998). In terms of sexual investment, some taxa display strikingly extensive variations in courting and mating strategies. This is the case of shorebirds (order Charadriiformes) in which all mating systems are encountered: from polyandry in phalaropes, dotterels or jacanas; to monogamy in oystercatchers and most lapwings; and polygyny in several sandpipers and plovers (Thomas et al., 2007). Getting a clear overview on mating systems and sex investments in different species and populations is of great importance, because it is likely these traits influence sex ratio and other demographic traits in species.

The knowledge of sex roles in shorebirds has been uneven, with information being usually available for Palearctic and Nearctic species, but mainly lacking in tropical species (Piersma et al., 1997; Kubelka, 2018). This is partially explained by the remoteness of those species, difficulty to monitor such populations in harsh environment or cryptic behavior and low population densities.

Lapwings (genus *Vanellus*) are mid-sized birds, with generally contrasted plumage and conspicuous behavior, making them easy to observe. They are distributed worldwide, expect in Antarctica and North America (del Hoyo et al., 2018). If some species have a well-described behavioral repertoire (e.g. Northern Lapwing *Vanellus vanellus*: Brown, 1926; Shrubb, 2010), some remain poorly described.

Seldom information are available on Andean Lapwing (*Vanellus resplendens*) general biology (Salvador, 1992; Capllonch & Quiroga, 2013; Simpson & Simpson, 2021). This is partially due to the distribution of the species in high elevations and sparsely-populated regions (Wiersma & Kirwan, 2020). Although anecdotical observations of nests and chick care were reported (Salvador, 1992; Capllonch & Quiroga, 2013), courtship behaviors of this species have not been described yet. To cover this knowledge gap, we reported observations of the courtship behavior and mating system of this High-Andes specialist, from a population on the Peruvian altiplano.

## Methods

### Study site

We observed Andean Lapwings on the shores of Laguna de Salinas in the Reserva Nacional de Salinas y Aguada Blanca, Arequipa province, Peru (16°21′36″S, 71°07′48″W, 4,315 m a.s.l.), between October 2021 and January 2022. Birds were observed on the North-West and South shores of the salina (Figure 1). The environments along the shores represented typical bofedales (Maldonado Fonkén, 2014), a peatland with cushion vegetation (likely *Distichia* sp.) and little puddles; as well as bare clay ground on the salina.

**Figure 1.**
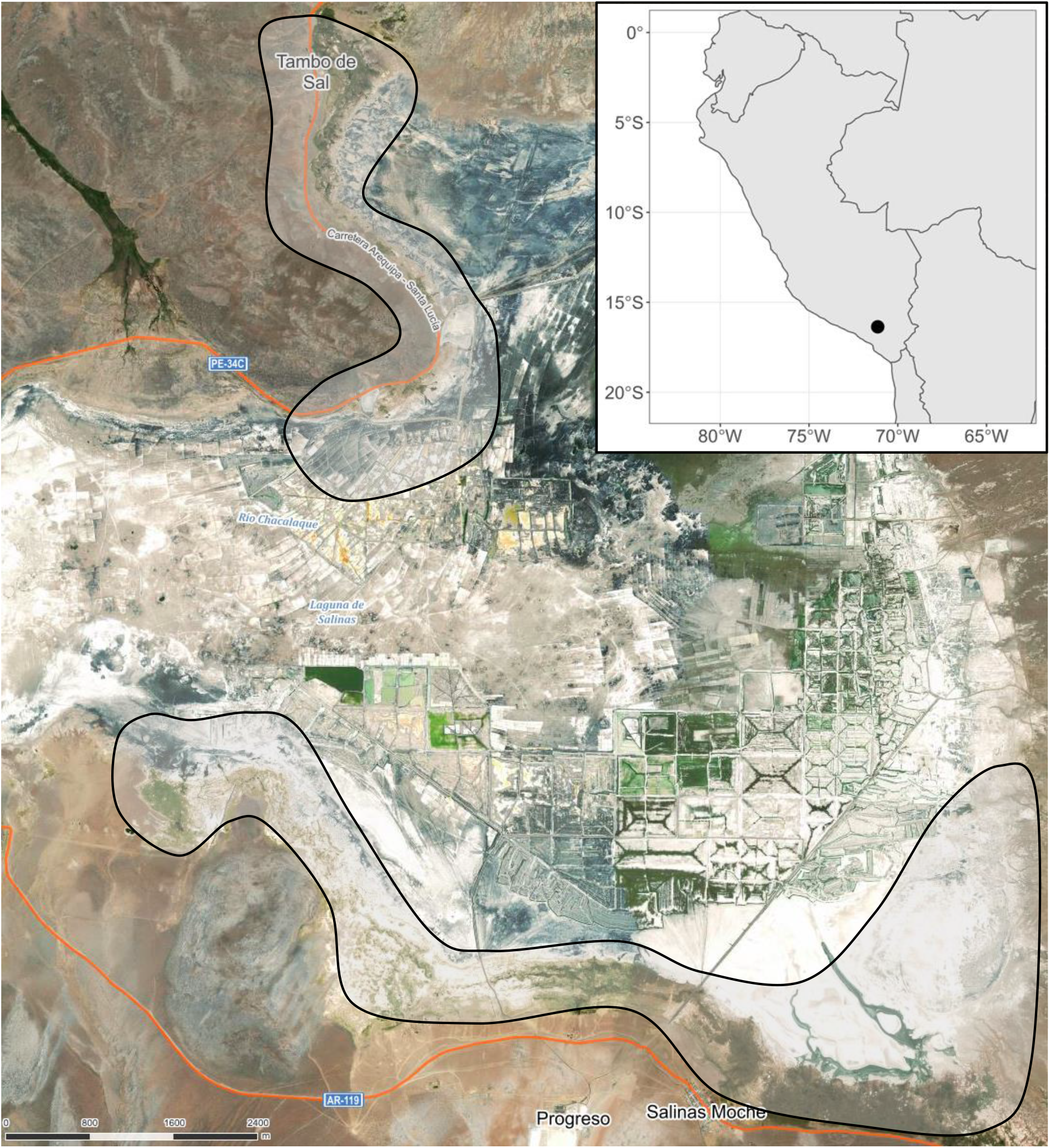
Location and aerial view from the study site (mapy.cz). Circled areas correspond to regularly monitored areas for lapwings. Central zone is mainly dedicated to mining activities and thus, not frequented by birds.

### Observations

We observed natural behavior of undisturbed birds, following established ÉLVONAL Shorebird Science protocol and ethogram for the observations (Székely & Kubelka, 2019). We usually restricted observations to the morning, between 0540 h and 1200 h, because of strong winds during the afternoon, making observations nearly impossible. Birds also seemed less active outside of this interval, mainly resting. We observed lapwings through scopes and binoculars (different models), from 50 to 400 m distance from focal birds, ensuring no disturbance. Concisely, we followed a pair and recorded the behavior of each individual every 20 s, for a period of 30 min. We allowed breaks when an individual wasn’t visible for a moment (maximum cumulating 30 min break). No evident dimorphism was observed between sexes, but individual coloration markings often allowed individual recognition. Sex was assigned only if copulation occurred. Birds were not marked, and we considered they belong to the same pair if they were observed a consecutive day in the same territory, within a distance of 300 m. Considering the low densities of lapwings at the time and their territorial behavior, we expect this method to be reliable.

In total, we recorded 20 pairs, between one and four times each from 21 November 2021 to 7 January 2022, representing a total of 49 observations. Pairs that didn’t display courtship activities were still included in the analysis, as we considered breeding season started when the first pairs displayed.

### Climate data

For each observation, we attributed temperature, wind speed and weather on the site. Temperature was given by local forecast on smartphones’ application. To estimate wind speed, we referred to Beaufort scale. Weather was split between “clear sky” if no cloud was visible, “cloudy” if sky blue was still visible between clouds, and “overcast” if sky blue wasn’t visible at all behind clouds. No observation happened during precipitations event.

### Statistical analysis

All statistical analyses were performed in R (v. 4.1.1) (R Core Team, 2021). To avoid misinterpretation of behavior, as stated by Maclean (1972), we gathered courting behavior and aggressive social behavior together. We considered the proportion of scans dedicated to courtship and territory defense behaviors, per observation, and averaged per each pair to avoid pseudo-replication. For sex investment analyses, only pairs which were safely sexed with copulation position were included (n=28). We tested sex investment and difference in courtship behaviors using unpaired two-samples Wilcoxon test.

## Results

### Timing and pair formation

Throughout the season, the number of observed individuals increased on the monitored sites, from at least 8 individuals during the first half of November, to a minimum of 38 in January. Small flocks of individuals were observed in early November, but never encountered later.

During the first time period of our field season, no courtship has been recorded, even though some pairs were already observed around, not necessarily on their definitive territories. First courting activity has been recorded on 21 November (Figure 2). This period corresponded with the first consistent rain precipitations, with first record on 20 November (Figure 2), later considered as the beginning of rain season. Anecdotal and sparce precipitations still occurred earlier in the season.

**Figure 2.**
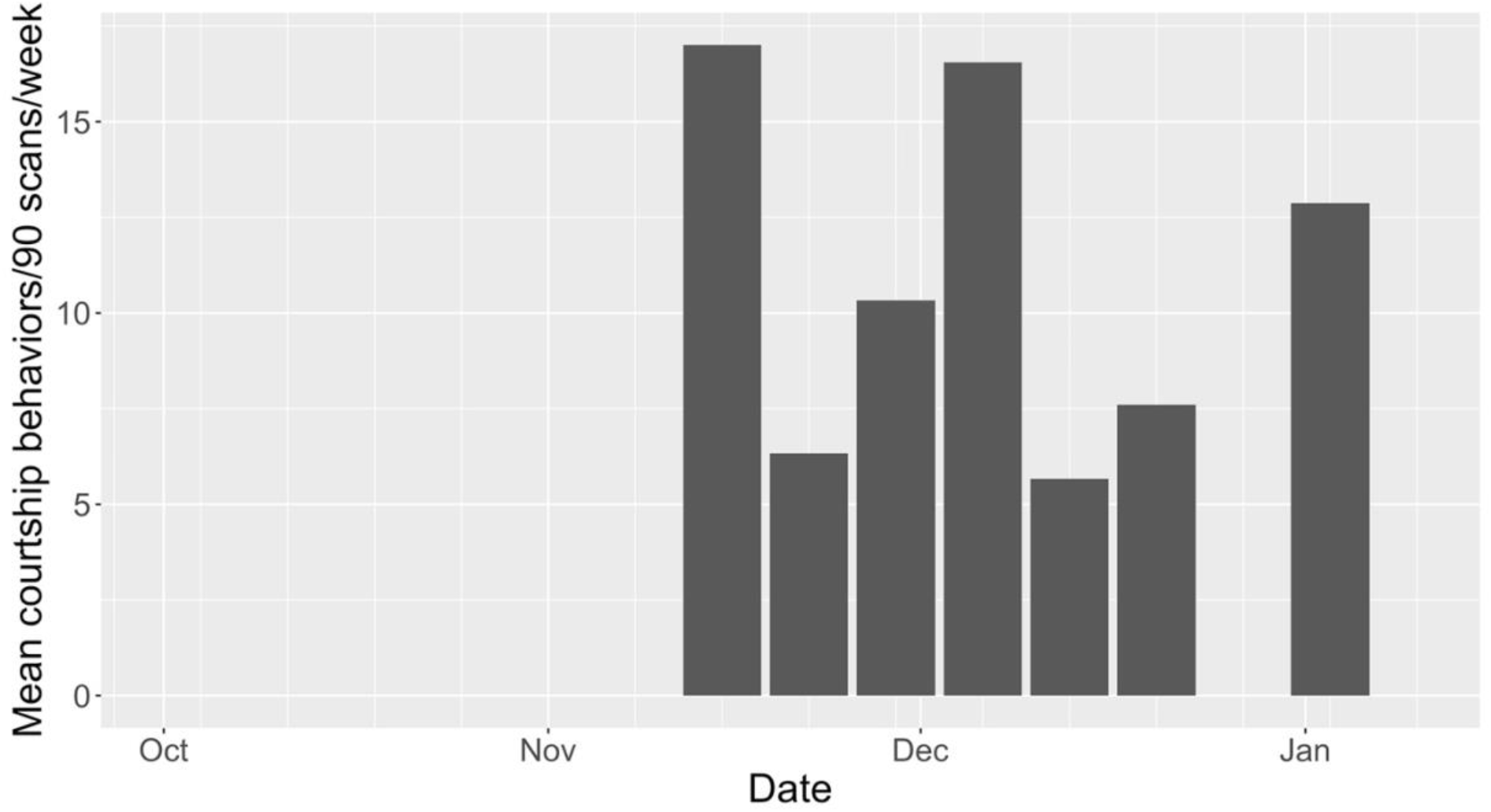
Weekly average number of courtship behaviors performed per observation session of 90 scans over the monitoring season. End October and the last week of December were not monitored. First rain precipitations were recorded on 30 Nov.

All observed pairs were formed by only one male and one female. No male has been seen copulating with more than one female. Out of the 20 pairs followed, only two did not performed mating behaviors during the observations and were not seen after. They were considered to not have established a territory yet.

### Courtship behavior

In terms of specific behaviors, lapwings performed limited aerial display. “Aerial display” was here described as flying back and forth above their territory, while emitting repeated screeching calls. During courtship, both partners get involved in an intimate display we referred as “choking”, as they lower their head, raise their tail and slightly open their wings several times, like a reverence. Calls are possibly emitted during this, but with distance we couldn’t attest it. We defined “ground display” as specific postures adopted while being on the ground: we identified the previous “reverence” posture sometimes displayed by only one individual, and an “about-to-sit” posture maintained for a few seconds. On one occasion, two individuals of the same pair staired at each other with straight up posture for a few seconds before copulating. This behavior was however coded as “head up”, so not courtship related. On one occasion, we observed a male walking with a flattened back and neck, coded as “flat running”, that we coded as courtship behavior.

Lapwings displayed the classical “scraping” sequence. Expected males pushed their breast into the ground and scraped to create a little depression. Usually females overtake this action, and both continue by pecking and throwing behind them small lining material towards the created scrape. On a few occasions, we witnessed the male sitting on the ground, and only starting scraping when the female came close.

No display appeared specific to pre- or post-copulation, and no particular call was heard, maybe due to distance. Before copulation, the female was observed crouching and extending her neck. In a few cases, the female adopted this position but the male didn’t answer by mounting her. In every successful copulation attempt, the male reached the female by walking, and never by landing on her.

Males significantly displayed more courtship behaviors, combining mating and territorial behaviors, than females (*w*=291, *P*=0.014) (Figure 3). When controlling for specific behaviors, males significantly chased more individuals out of their territory (*w*=27, *P*=0.074), and invested more time on scraping (*w*=156, *P*=0.039) (Figure 4).

**Figure 3.**
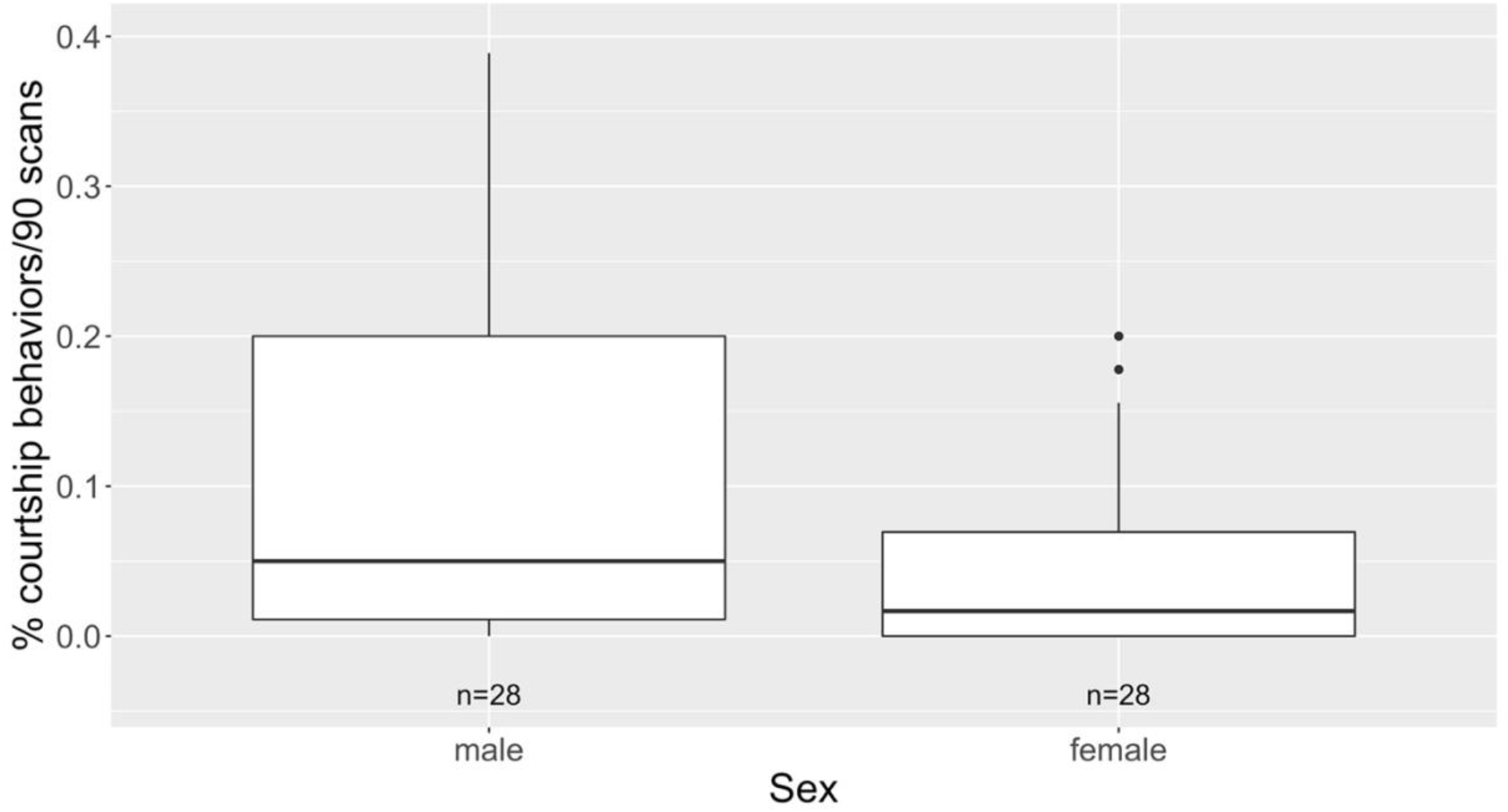
Percentage of scans allocated to courtship behaviors per observation session of 90 scans according to sex. Here, only observations during which sex identity was confirmed by copulation are included.

**Figure 4.**
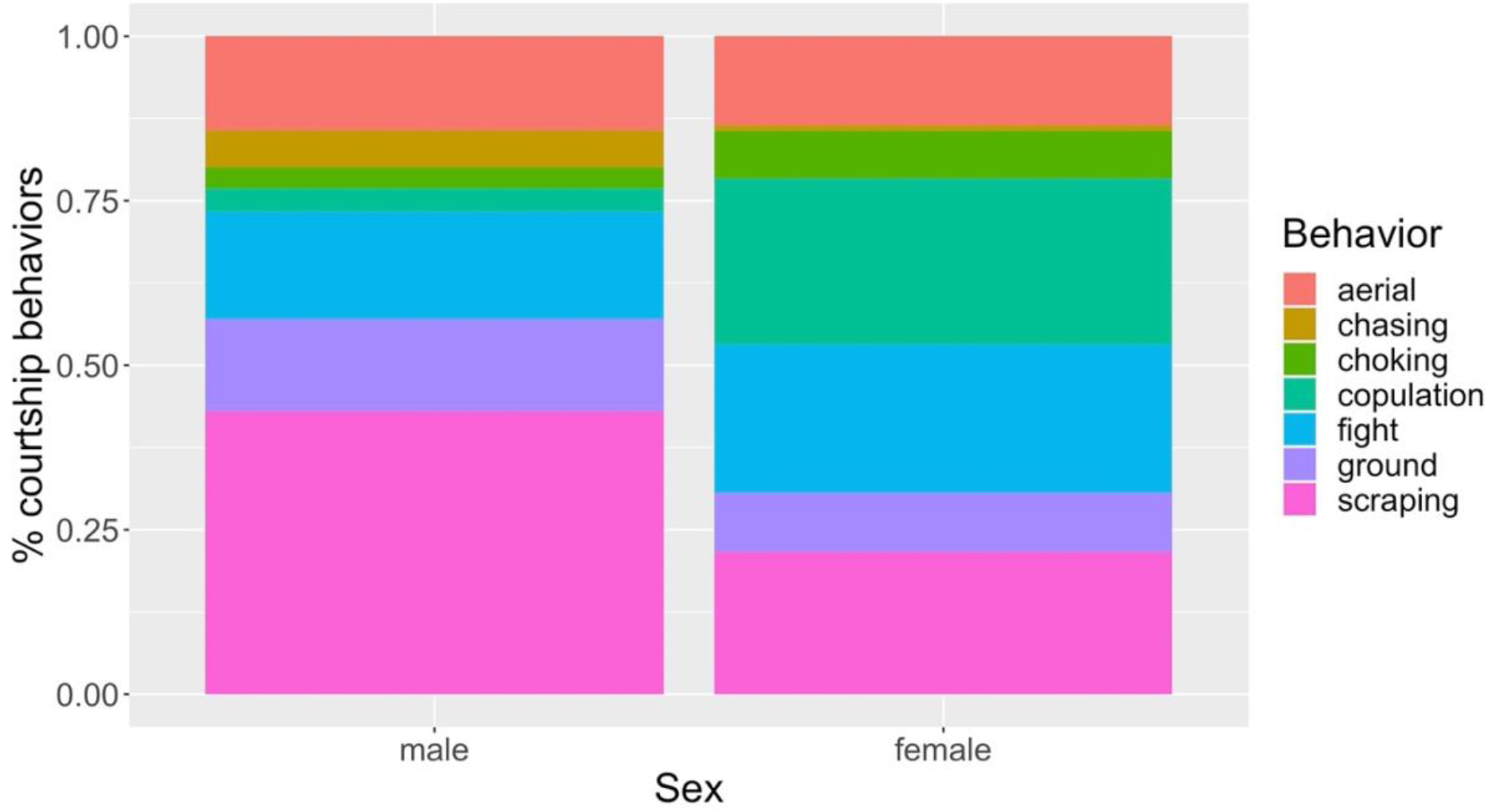
Percentage of displayed courtship behaviors split by categories according to sex. Here, only observations during which sex identity was confirmed by copulation are included (n=28). “Aerial” includes aerial display; “chasing” and “fight” includes intra- and interspecific interactions, “copulation” includes also pre-copulation for females, “ground” includes reverence and flat-run posture, while “choking” refers to intimate display when both individuals participated, “scraping” includes all behaviors linked with scraping sequence.

Although our data were skewed towards early morning conditions with low wind and cold temperatures, we found no significant effect of both wind strength, weather or temperature on courting activity intensity (all *P*>0.05).

### Territorial behavior

Both partners were potentially invested in territory defense: aggressive behaviors can be displayed as the “extremely head up” raised posture, or the previously described “reverence” posture towards the opponent. When two conspecific pairs met, aggressive behaviors were often performed by the two partners. On one occasion, both partners managed to chase conspecifics, and started the “reverence” display as a what-could-be-anthropocenteredly-called celebration. Aggressive behaviors were also displayed towards heterospecific: several passerine species (e.g., Andean Negrito *Lessonia oreas*, Common Miner *Geositta cunicularia*), other shorebirds (e.g., Gray-breasted Seedsnipe *Thinocorus orbignyianus*), Andean Gull (*Chroicocephalus serranus*), Andean Goose (*Oressochen melanopterus*), James’s Flamingo (*Phoenicoparrus jamesi*), human observers, or llama (*Llama glama*) and alpacas (*Vicugna pacos*). In these cases, lapwings performed aerial display towards bigger species, the latter often not reacting, or aggressive chasing resulting in flight of the smaller species.

### Nesting behavior

We observed at least two scrapes being built by lapwings. Scrapes were little depression built on cushion vegetation, with little vegetation as lining around it. Despite some pairs starting to court early, we observed no laid egg after one month and half, and these pairs were still actively courting and building new scrapes by early January.

## Discussion

As the population of lapwings increased throughout the season, it is supposed that new individuals were joining the site at this period. It is then likely that Laguna de Salinas represented a breeding site for this species, and birds wintering elsewhere are returning here. Andean Lapwing are known to perform an altitudinal migration and spend non-breeding season at lower elevations (Wiersma & Kirwan, 2020).

Lapwings only displayed courtship behaviors during the second part of our field season, from late November onwards. This seems to indicate a certain seasonality within this population regarding breeding timing, contrary to other tropical populations that breed all year long (Goymann & Helm, 2014). As lapwings started to court as the rain precipitations became more frequent, rainfalls seem to positively influence breeding season’s start in this species. Indeed, rainfall are expected to represent a possible circannual zeitgeber in tropical regions (Goymann & Helm, 2014). Here, the rainfall could have either acted as an indicator for individuals to start breeding, or more likely move to their breeding grounds.

Our population seemed to display social monogamy, based on strong territoriality and absence of observation of individuals mating several partners. This is consistent with breeding strategies in majority of *Vanellus* species, excepted Northern Lapwing in which some males are polygynous (Shrubb, 2010). Even non-territorial individuals were observed in pairs, suggesting either strong pair bonding during the year, or pair formation prior to breeding season. Both mating strategies have been described in the Northern Lapwing (Brown, 1926).

The only two pairs observed not courting during the breeding season may suggest that territory defense and breeding activity start soon after settlement of birds on their breeding ground.

Overall, the number of displays performed during courtship appeared rather limited. According to Maclean (1972), it is expected that several displays appear similar between aggression and courting, and we could not exclude that we categorized incorrectly certain behaviors. For instance, birds of the same pair stared at each other in a “head up” position, usually associated with aggressiveness even in this species, before copulating.

Maclean (1972) insisted on the fact that *Vanellus* genus is characterized by post-copulatory displays including raised wings. Several other lapwing species seem to perform wing displays or calls before, during or after mounting (Hall, 1964, 1965; Little, 1967; Muralidhar & Barve, 2013). This was not noticed in our population.

Wing patterns are generally highly contrasted in *Vanellus* species, and they are expected to be shown during displays, either aggressive or affiliative (Maclean, 1972; Shrubb, 2010). Out of post-copulatory displays, lapwings show their wings during elaborate aerial display. This includes, flying back and forth and suddenly up and down, or abruptly switching wing position on a 45° axis to make pattern visible from different directions (Brown, 1926; Skead, 1955; Little, 1967; Ward, 1989; Saxena & Saxena, 2013). Compared to these observations in Old World species, aerial display in Andean Lapwing was rather timid, without abrupt turns or extensive wing showing while flying.

The described “reverence” posture has already been reported in other *Vanellus* species, in both territorial (Brown, 1926; Hall, 1964; Walters & Walters, 1980) and mating contexts (Little, 1967). This probably also refers to the “bowing” display described, and found in other more distant species (e.g. Snowy Sheathbill *Chionis albus*: Jones, 1963). Little (1967) argued for the fact that this behavior usually resulted in copulation, but this was not the case in our population. The scraping and nest building sequence observed here was similar to the ones described for other congenerics (Hall, 1964, 1965; Little, 1967; Kumar, 2015). We may only point out that on several occasions, the supposed male just sat on the ground and waited for the female to come closer to initiate scraping. Oppositely in other studies, scraping seems to attract the female (e.g. Shrubb, 2010).

Contrary to congenerics, we never observed a fly-by copulation, where the male engaged in copulation landing directly on or next to the female (e.g. Brown, 1926).

It seemed surprising that we never observed birds in egg laying phase, despite some pairs observed courting quite early. Indeed, Argentinian populations were observed laying eggs within a month after territory establishment (Capllonch & Quiroga, 2013), although nesting period seems widely spread until February (Salvador, 1992; Wiersma & Kirwan, 2020). We are unable to tell if eventual eggs were laid and quickly destroyed or depredated, or if birds had a delayed egg laying and incubation. It seems rather unlikely that clutches were predated, as artificial nests in the area experienced extremely low predation rates (AR, 2022, pers. comm.). However, llamas, alpacas and sheep, present in high densities in breeding areas, may trample on nests, but we found no broken eggs either. Hall (1964; 1965) noted strong displays when cattle approached a nest, so it is possible that the pair reactive to it actually had nests. In locations with no cattle, we still failed to find nests.

Some may argue birds were still on their non-breeding grounds while displaying pair bonding. Other lapwings have been observed courting and scraping prior to the breeding season, later arriving in pairs on their breeding grounds (Brown, 1926; Hall, 1964). Andean Lapwing is supposed to be rather sedentary, with eventual altitudinal migration (Wiersma & Kirwan, 2020). However, number of observed individuals increased during the season, indicating individuals were rather settling in the area during this period, and the site corresponded to a breeding ground.

Taken together, Andean Lapwings seem to share many behaviors with their *Vanellus* congenerics, but set themselves apart to display limited number of behaviors during breeding season. We still miss information on other high-elevation-breeding species to validate such assumptions. To complete the behavioral portrait of these species, additional observations on incubation, brood care and non-breeding periods, and data on reproductive investments are required.

## Acknowledgments

We thank the team of Sernanp for their authorization to work within the reserve. The fieldwork expedition was covered by ÉLVONAL Shorebird Science grant.

## Notes

### Competing Interest Statement

The authors have declared no competing interest.

